# Metabolic dysfunction and metformin modify plasma extracellular vesicle signaling to prostate cancer cells

**DOI:** 10.64898/2025.12.07.692844

**Authors:** Michael Seen, Pablo Llevenes, Heejoo Kang, Yuhan Qiu, Christina S. Ennis, Andrew Chen, Sara Alexanian, Devin Steenkamp, Stefano Monti, Gerald V. Denis

**Affiliations:** Cancer Center, Boston University Chobanian and Avedisian School of Medicine, Boston, Massachusetts; Department of Medicine, Computational Biomedicine Section, Boston University Chobanian and Avedisian School of Medicine, Boston, Massachusetts; Bioinformatics Program, Boston University, Boston, Massachusetts; Sara Alexanian, Associate Medical Director, Global Drug Safety, Biogen, Cambridge, Massachusetts; Section of Endocrinology, Diabetes, Nutrition and Weight Management, Boston University Chobanian and Avedisian School of Medicine, Boston, Massachusetts; Department of Pharmacology and Experimental Therapeutics, Boston University Chobanian and Avedisian School of Medicine, Boston, Massachusetts; Section of Hematology/Oncology, Boston University Medical Center, Boston, Massachusetts

**Keywords:** prostate cancer, type 2 diabetes, metformin, extracellular vesicles, EV-derived RNA, microRNA

## Abstract

Type 2 diabetes (T2D) is associated with altered outcomes in established prostate cancer, but the underlying mechanisms remain unclear. We examined whether whole-body metabolic state and systemic metformin treatment are associated with changes in circulating plasma extracellular vesicle (EV) signaling to prostate cancer cells. Plasma EV-derived miRNAs were profiled in six T2D and six non-diabetic (ND) donors, selected mimics were tested in DU145 cells for effects on transcription of *ZEB1*, a master regulator of tumor progression. Recipient cell transcriptional responses to human EV-enriched fractions were also assessed by RNA sequencing using six T2D and two ND donors. DU145 cells were also transfected with plasma EV-derived RNA from high fat diet (HFD)-fed mice with or without metformin treatment. Human miRNA profiling identified miR-192-5p, miR-106b-5p, and miR-29b-3p as candidates with greater than two-fold elevation and nominal *P*<0.05; none of the 84 assayed miRNAs remained significant after multiple-testing correction. Synthetic miR-192-5p and miR-106b-3p mimics increased *ZEB1*, whereas miR-29b-3p had no effect on *ZEB1*. T2D donor EV exposure was associated with broad range of aggressive responses, including cytoskeletal, extracellular matrix, mitotic, and RHO-GTPase-related pathways. Mouse EV-derived RNA from metformin-treated HFD mice induced lower DU145 migration than RNA from untreated HFD mice (12.23% versus 20.25% mean membrane coverage; n=3 mice/group; Welch *P*=0.0246). Matrigel invasion analysis showed less invasion for HFD+metformin than HFD alone (4.89% versus 7.89%; n=3 mice/group; Welch *P*=0.0302). These findings support a strong association between abnormal systemic metabolism and circulating EV signaling to drive prostate cancer progression, together with a metformin-driven reduction in the tumor progression activity of plasma EV-derived RNA. The responsible RNA species and molecular mechanisms will be central to future research.

## Introduction

Obesity and its metabolic complications have long been associated with increased cancer incidence, progression, and cancer-related mortality [1–3]. Type 2 diabetes (T2D) is a prevalent, chronic metabolic disorder characterized by hyperglycemia and systemic metabolic dysfunction [4]. Global diabetes prevalence has risen markedly over recent decades [5] and contemporary epidemiologic analyses increasingly emphasize that obesity and diabetes drive not only elevated cancer risk but also worse prognosis after a cancer diagnosis [6].

In prostate cancer, the relationship with T2D is complex. Diabetes has been associated with lower incidence in some populations, particularly early-stage disease, with associations varying by stage and study design [7–9]. After diagnosis, diabetes and poor glucose control have been associated with higher mortality or worse outcomes in some clinical settings [10–15]. However, a large T2D cohort reported lower prostate cancer-specific mortality at higher glycated hemoglobin (HbA1c) levels [16], and treatment-related survival differences varied by body mass index (BMI) and glycemic status [17]. Cardiometabolic disease is also associated with increased risk for developing aggressive prostate cancer [18], a distinct endpoint from post-diagnosis progression. Associations with biochemical recurrence remain inconsistent [19]. These findings motivate our novel investigation of how systemic metabolic state alters established prostate cancer biology.

Circulating extracellular vesicles (EVs) represent one potential mechanism through which systemic metabolic dysfunction reprograms tumor behavior. EVs are membrane-bound particles released by cells and can carry proteins, lipids, and nucleic acids, including microRNAs (miRNAs); current field guidance emphasizes careful operational definition of EV preparations and avoidance of unsupported claims about vesicle subtype or biogenesis [20]. Reviews of prostate cancer and obesity-related signaling have highlighted EVs and EV-derived miRNAs as plausible mediators of tumor-microenvironment and systemic communication [21,22]. Prior published work from our group has linked metabolically altered plasma- or adipocyte-derived EVs with cancer aggressiveness in prostate and breast cancer models, including epithelial-mesenchymal transition (EMT)-related signaling, cytoskeletal remodeling, migration, organoid aggressiveness, tumor immune tolerance, and pro-metastatic behavior [23–27]. Together, these studies provide a potent rationale for examining whether metabolic status, particularly T2D, is accompanied by changes in circulating EV-derived signals that modify prostate cancer-cell responses to more aggressive phenotypes.

In this study, we investigated whether systemic metabolic state and metformin treatment are associated with altered plasma EV signaling to human prostate cancer cell models. We first compared EV-derived miRNA profiles from T2D and ND donors and tested selected candidate miRNAs for effects on the EMT-associated transcription factor ZEB1. We then examined the global transcriptional response of DU145 cells exposed to intact human plasma EV-enriched fractions. Because medication exposure in the human cohort was observational and potentially confounded, we used HFD-fed mice with or without metformin treatment to test whether systemic metformin modifies the functional activity of plasma EV-derived RNA, using prostate tumor cells as the readout. We hypothesized that metabolic dysfunction would be associated with altered plasma EV signaling and that metformin treatment under controlled metabolic conditions would attenuate pro-migratory activity carried by plasma EV-derived RNA.

## Materials and Methods

### Cell culture

DU145 human prostate cancer cells were obtained from the American Type Culture Collection (ATCC; HTB-81) and maintained in Dulbecco’s Modified Eagle Medium (DMEM; Gibco, 11995073) supplemented with 10% fetal bovine serum (Corning, 35016CV) and 1% penicillin-streptomycin (Gibco, 15140122). Cells were cultured at 37°C in a humidified atmosphere containing 5% CO□. Cells were confirmed negative for *Mycoplasma* contamination using the MycoStrip test kit (InvivoGen, rep-mys-50) before experimental use and were used within 10 passages.

### Human plasma donors and blood processing

Participants with T2D were recruited through the Boston Medical Center Diabetes Outpatient Clinic under a Boston University Institutional Review Board-approved protocol. T2D status was defined by an HbA1c value >6.5%. Individuals with active cancer were excluded from the T2D cohort. Whole blood was collected by certified phlebotomists into tubes containing Acid-Citrate-Dextrose Solution B (Vacutainer, 364816). Blood processing was performed at room temperature and completed within 1 h of collection. Plasma was separated by centrifugation for 20 min at 200 × *g*, transferred to a fresh tube, and depleted of platelets by two sequential centrifugation steps at 2,500 × *g* for 15 min, with transfer to a fresh tube after each step. Platelet-depleted plasma was aliquoted and stored at -80°C until EV isolation.

Non-diabetic (ND) plasma samples included study-associated donors and commercially sourced donors from Research Blood Components. Commercial records designated these donors as non-diabetic and provided limited demographic information; HbA1c, detailed metabolic measurements, comorbidity history, medication history, and complete cancer history were not available. Samples were not pooled. Plasma EV miRNA profiling included six T2D and six ND donors, whereas recipient-cell RNA sequencing included six T2D and two ND donors. Donor demographic and clinical characteristics are listed in Table 1; documented medications and experimental use are listed in Table 2.

**Table 1.** Characteristics of six T2D and six ND donors used for plasma EV-derived miRNA profiling. . Recipient-cell RNA sequencing used the six T2D donors and ND donors 1009 and 1014 (Table 2). Donor preparations were not pooled. T2D, type 2 diabetes; ND, non-diabetic; BMC, Boston Medical Center; BMI, body mass index; HbA1c, glycated hemoglobin; M, male; F, female; NR, not reported or available; y, years; -, not applicable. Commercial ND donors were designated non-diabetic by the vendor; HbA1c, detailed metabolic testing, comorbidity history, and complete cancer history were unavailable.

| **Study ID** | **Group / source** | **Age** | **Sex** | **Ethnicity** | **BMI** | **HbA1c (%)** | **Diabetes duration** |
|---|---|---|---|---|---|---|---|
| 1007 | T2D / BMC | 41 | M | Black | 34.6 | 6.8 | <5 y |
| 1008 | T2D / BMC | 48 | M | Japanese descent; Peruvian background | 28.71 | 8.4 | >10 y |
| 1010 | T2D / BMC | 60 | F | Black | 37.04 | 8.4 | >10 y |
| 1011 | T2D / BMC | 56 | M | White | NR | 7.1 | >10 y |
| 1012 | T2D / BMC | 60 | M | Black | NR | NR | NR |
| 1013 | T2D / BMC | NR | F | NR | NR | NR | NR |
| 1009 | ND / study-associated | NR | M | White | NR | NR | - |
| 1014 | ND / study-associated | NR | M | White | NR | NR | - |
| 1015 | ND / commercial | 54 | M | White | NR | NR | - |
| 1016 | ND / commercial | 62 | M | Black | NR | NR | - |
| 1017 | ND / commercial | 56 | M | White | NR | NR | - |
| 1019 | ND / commercial | 46 | M | White | NR | NR | - |

**Table 2.** Medication histories and experimental inclusion are shown for the same donors as in. Table 1. miRNA qPCR denotes plasma EV-derived miRNA profiling (6 T2D and 6 ND donors); RNA-seq denotes recipient-cell RNA sequencing after donor EV treatment (6 T2D and 2 ND donors). Donor preparations were not pooled. Medication names were standardized from the donor source records. “Not listed” means metformin was absent from the available medication list and does not establish non-use; NR denotes information not reported or available. Missing medication entries were not inferred. T2D, type 2 diabetes; ND, non-diabetic; EV, extracellular vesicle; qPCR, quantitative polymerase chain reaction; RNA-seq, RNA sequencing.

| Study ID | Metformin | Other documented medications | miRNA qPCR | RNA-seq |
|---|---|---|---|---|
| 1007 | Yes | Dulaglutide | Yes | Yes |
| 1008 | Yes | Dulaglutide; empagliflozin | Yes | Yes |
| 1010 | Not listed | Insulin; dulaglutide; empagliflozin | Yes | Yes |
| 1011 | Yes | Empagliflozin; insulin glargine; semaglutide | Yes | Yes |
| 1012 | Not listed | Insulin glargine; insulin aspart; dulaglutide; empagliflozin; atorvastatin; lisinopril | Yes | Yes |
| 1013 | Yes | Atorvastatin; dulaglutide; lisinopril | Yes | Yes |
| 1009 | NR | NR | Yes | Yes |
| 1014 | NR | Lisinopril; statin | Yes | Yes |
| 1015 | NR | NR | Yes | No |
| 1016 | NR | NR | Yes | No |
| 1017 | NR | NR | Yes | No |
| 1019 | NR | NR | Yes | No |

### Mouse blood collection and plasma processing

Mice were anesthetized with isoflurane, and adequate anesthesia was confirmed by absence of a toe-pinch reflex. Blood was collected by retro-orbital sampling using EDTA-coated capillary and collection tubes. Plasma was separated by centrifugation for 20 min at 200 × *g*, followed by two sequential centrifugation steps at 2,500 × *g* for 15 min, with transfer to a fresh tube after each step. Platelet-depleted plasma was aliquoted and stored at -80°C.

### Plasma EV-enriched fraction isolation and human nanoparticle tracking analysis

EV-enriched fractions were isolated from platelet-depleted plasma by size-exclusion chromatography using an automated fraction collector (Izon, AFC-V2) and qEV Single Gen 2 columns with a nominal pore size of 35 nm (Izon, ICS-35). The elution buffer consisted of calcium- and magnesium-free DPBS (Cytiva, SH3002801) supplemented with 1 mM EDTA (Invitrogen, 15575020) and filtered through a 0.2-µm membrane (Fisher, FB12566500) on the day of isolation. For each preparation, 150 µL of platelet-depleted plasma was loaded onto the column. The first 0.87 mL of eluate was discarded, and the subsequent 0.75 mL was collected as the EV-enriched fraction. Fractions were stored at 4°C and used within 24 h.

Particle-size distributions and concentrations were measured by nanoparticle tracking analysis (NTA) using a NanoSight NS300 (Malvern Panalytical) to characterize the human EV-enriched preparations [20]. Human donor measurements are presented in Supplementary Figure S1 and Supplementary Table S1. For experiments with human-derived EV-enriched fractions, treatment doses were normalized to 100,000 particles per DU145 cell using the concentrations measured by NTA.

### Gene-expression RT-qPCR

Total RNA was isolated from DU145 cells using the RNeasy Mini Kit (Qiagen, 74104) with on-column DNase digestion (Qiagen, 79254). A total of 500 ng RNA was reverse transcribed using the QuantiTect Reverse Transcription Kit (Qiagen, 205311). Quantitative PCR was performed using TaqMan Gene Expression Master Mix (Applied Biosystems, 4369016) with 25 ng cDNA per reaction. Human gene-expression assays were ACTB (Hs00357333_g1) and ZEB1 (Hs01566408_m1).

### Plasma EV miRNA profiling

RNA was isolated from plasma EV-enriched fractions using the miRNeasy Serum/Plasma Kit (Qiagen, 217184). First-strand synthesis was performed using the miRCURY LNA RT Kit (Qiagen, 339340). The Human Cancer Focus miRCURY LNA miRNA PCR Panel (YAHS-102Y; Qiagen, 339325), together with miRCURY LNA SYBR Green PCR chemistry (Qiagen, 339345), was used to profile 84 cancer-associated miRNAs and assay controls. Six independent T2D donors and six independent ND donors were analyzed. Ct values were processed using Qiagen GeneGlobe. The original candidate-selection workflow identified miRNAs showing >2-fold elevation with nominal *P*<0.05. For multiple-testing assessment, nominal *P* values across all 84 assayed miRNAs were adjusted using the Benjamini-Hochberg procedure [28], with *q*<0.05 used as the FDR significance threshold.

### RNA sequencing and bioinformatic analysis

DU145 cells were treated for three days with intact plasma EV-enriched fractions derived from six T2D donors and two ND donors, followed by RNA sequencing. Untreated DU145 recipient cell samples were also sequenced as a control condition for the descriptive Principal Components Analysis (PCA) and the T2D versus untreated control comparison. For selected donor preparations, multiple recipient cell treatment wells were sequenced; these were within-donor recipient cell replicates and did not constitute additional independent human donors. Total RNA was submitted to Novogene for sequencing. Sequencing reads were processed using the nf-core/rnaseq workflow [29] with the human GRCh38/hg38 reference genome. Gene level count matrices were analyzed using DESeq2 [30]. The biological donor comparison therefore comprised six independent T2D donors and two independent ND donors, with repeated recipient cell wells retained as within-donor replicates rather than counted as additional human subjects.

For the T2D versus ND volcano plot analysis, genes were classified as differentially expressed using adjusted *P*<0.05 and an absolute log2 fold change >1. PCA was used descriptively to visualize global transcriptional patterns and was not used to test medication-specific effects. For the Figure 2C heatmap, genes were restricted to those meeting adjusted *P*<0.05 and absolute log2 fold change >1 in both the T2D versus ND and T2D versus untreated control comparisons.

**Figure 1.**
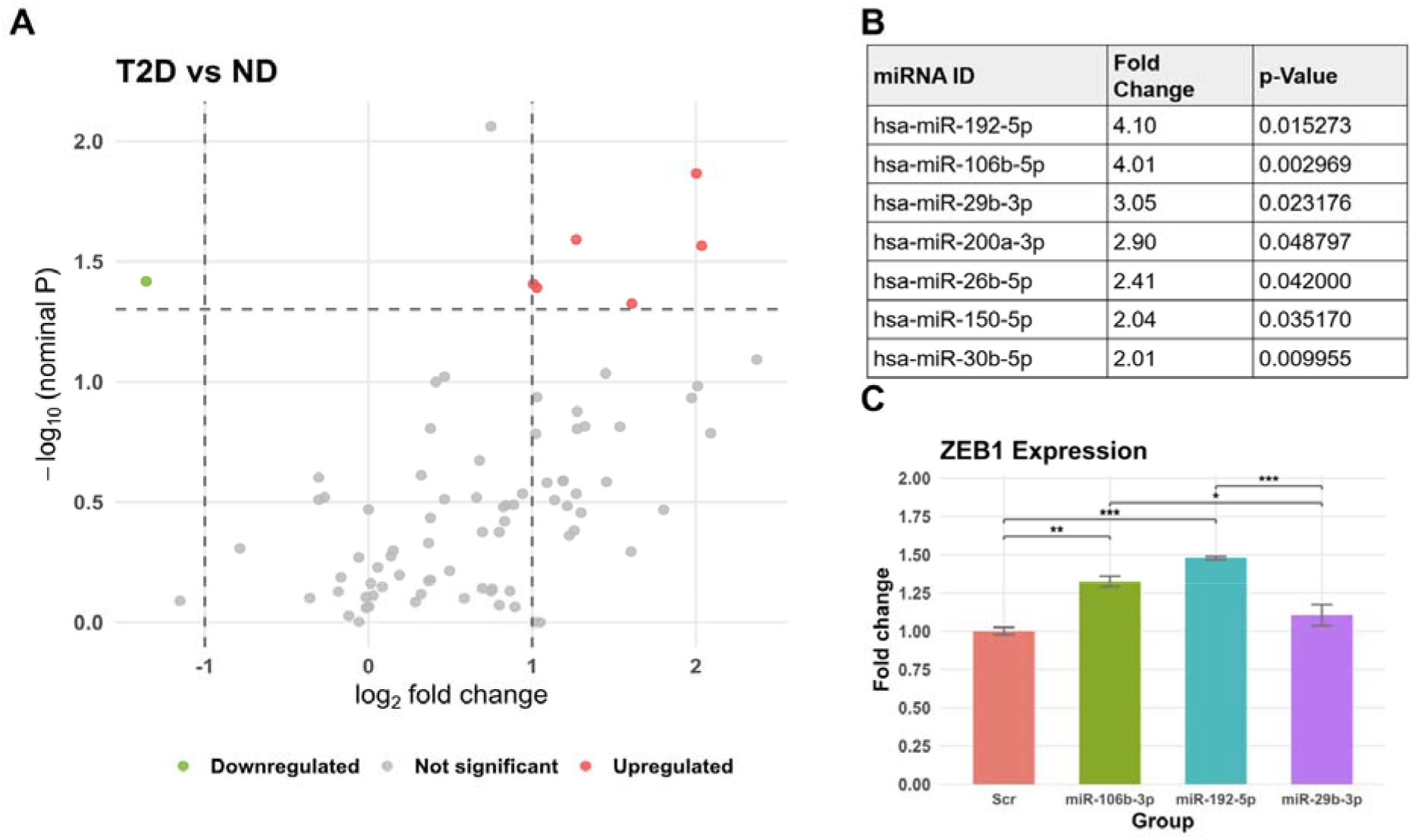
**Plasma EV miRNA profiling in T2D and non-diabetic donors.**A. qPCR-array analysis of cancer-associated miRNAs in plasma EV-derived RNA from T2D and ND donors (n=6 donors/group). Screening criteria were nominal *P*<0.05 and fold change >2. B. Candidate miRNAs meeting nominal screening criteria. Screening *P* values in panels A-B are nominal; none of the 84 assayed miRNAs remained significant after Benjamini-Hochberg correction (q<0.05). C. *ZEB1* expression after synthetic miRNA mimic transfection. Data are mean ± SEM (n=3 samples/condition); one-way ANOVA followed by Tukey’s multiple-comparisons test. Symbols indicate Tukey-adjusted pairwise comparisons: \**P*<0.05, \*\**P*<0.01, \*\*\**P*<0.001.

**Figure 2.**
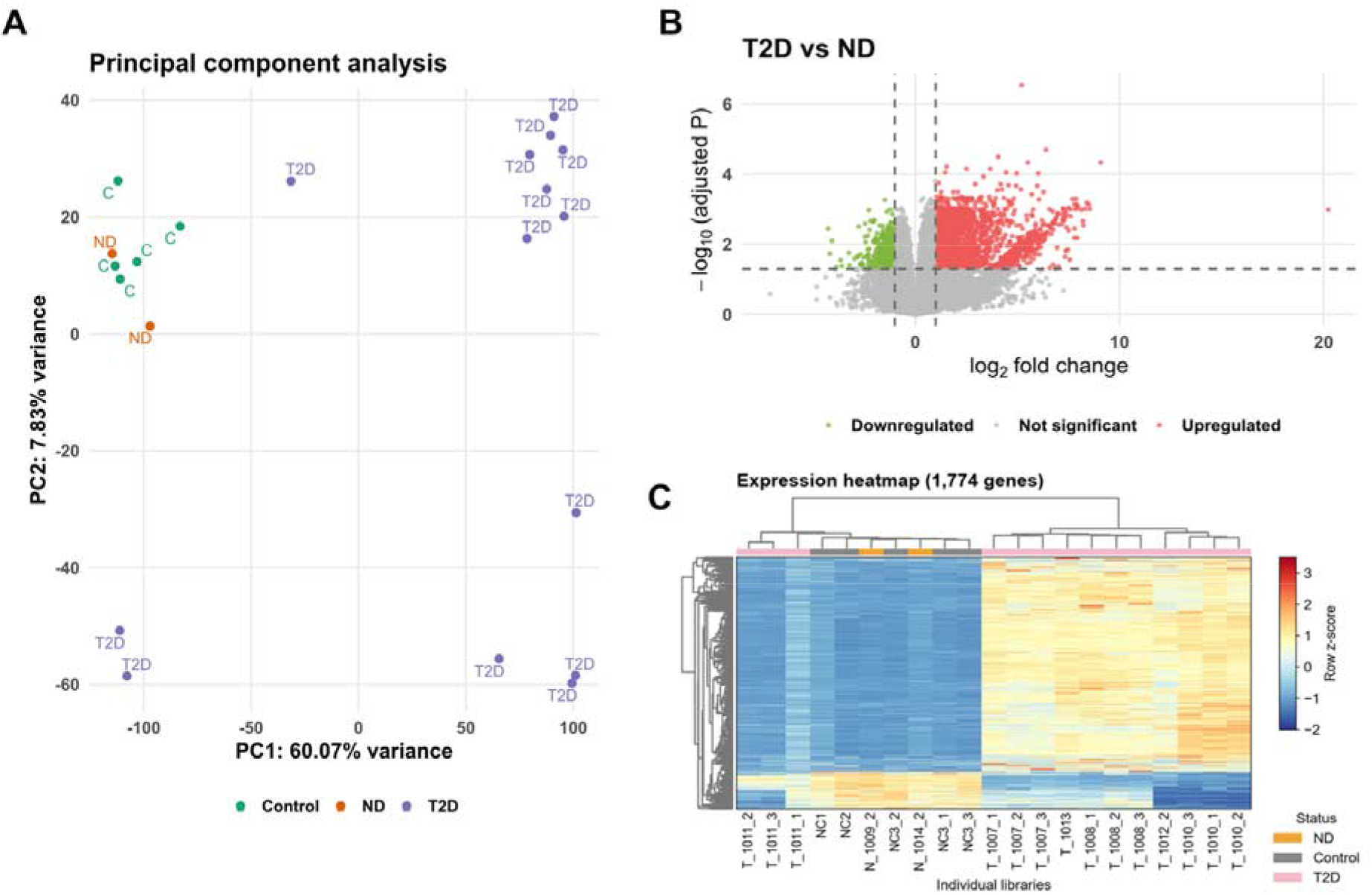
Transcriptional responses of DU145 cells to plasma EV-enriched fractions from T2D and ND donors. The human RNA-sequencing cohort comprised six independent T2D donors and two independent ND donors; repeated sequenced recipient-cell wells from the same donor were within-donor replicates and did not increase the independent human donor n. Untreated DU145 recipient-cell samples were included as a control condition. A. PCA of recipient-cell RNA-sequencing samples. B. T2D versus ND differential expression analysis using adjusted *P*<0.05 and |log2FC|>1. C. Heatmap generated from the more restrictive gene set meeting these criteria in both the T2D versus ND and T2D versus untreated control comparisons.

For the T2D-versus-ND comparison in Figure 3B–C, gene-level estimated counts were summed across technical libraries from the same donor and rounded to integers, yielding one count profile per donor. The analysis included six T2D and two ND donors; untreated samples were excluded. Genes with at least 10 counts in at least two donors were retained, yielding 20,030 genes. Differential expression was calculated using PyDESeq2 v0.5.4 [31], with condition as the design variable (∼condition) and a Wald test comparing T2D with ND. Standard procedures were used for size-factor normalization, dispersion estimation, and assessment of outliers using Cook’s distance. Figure 2 presents the original archived PCA and differential-expression results.

**Figure 3.**
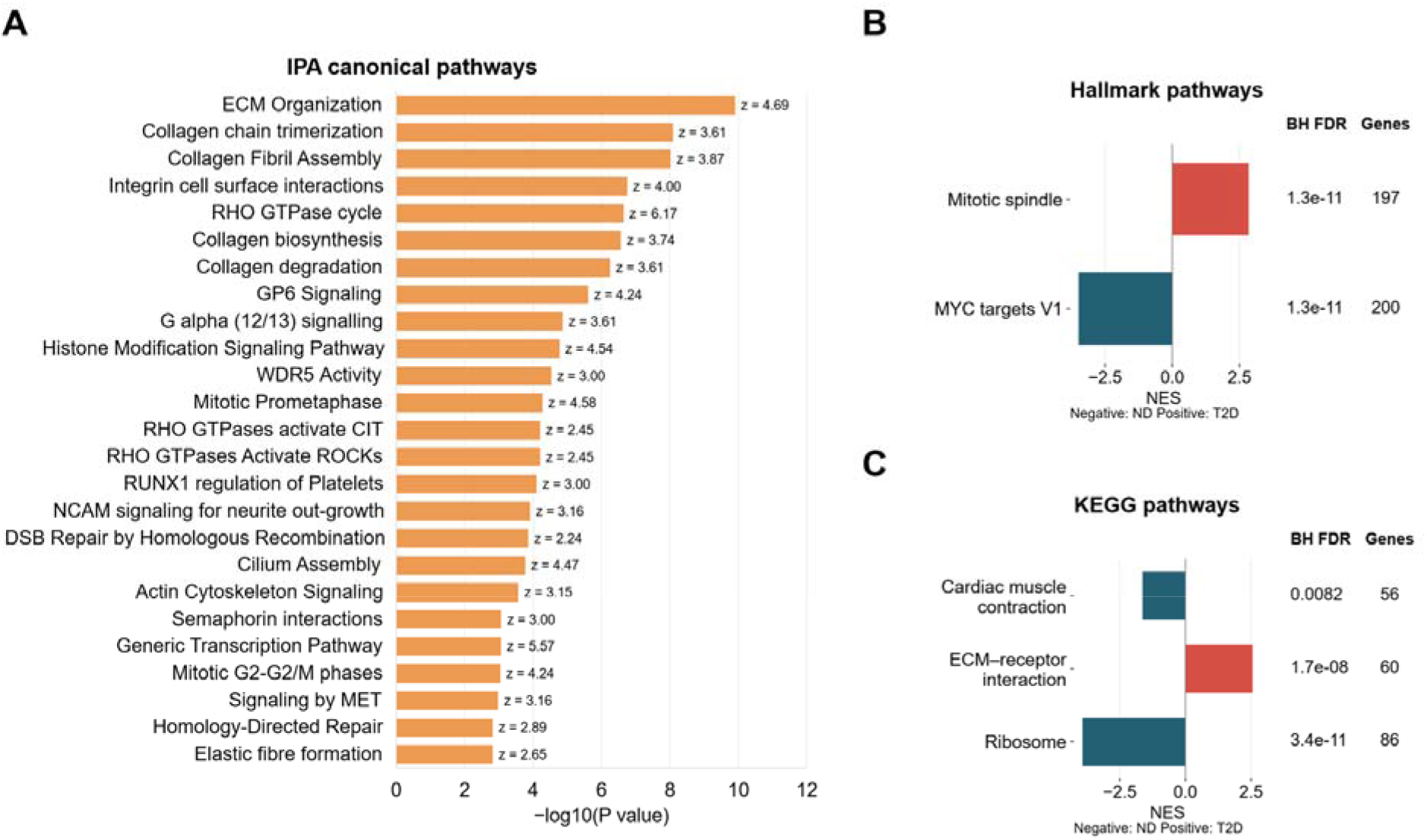
Pathway analyses of transcriptional responses in plasma EV-treated DU145 cells. A. Archived IPA canonical pathway enrichment; bar length represents -log10(*P* value), and labels at the bar ends report the archived activation z-score, a computational prediction rather than a direct measurement of pathway activity. B-C. Newly recomputed donor-level pre-ranked enrichment from Batch 2 counts, with technical libraries aggregated within each of six T2D and two ND donors. B. Hallmark mitotic spindle and MYC targets V1. C. KEGG cardiac muscle contraction, ECM receptor interaction and ribosome. Positive NES indicates enrichment toward T2D; negative NES indicates enrichment toward ND. BH FDR reports Benjamini-Hochberg-adjusted nominal enrichment *P* values within each full tested collection. Genes indicates gene-set membership in the ranked universe.

### Pathway analysis

Pathway analyses included Ingenuity Pathway Analysis (IPA; QIAGEN), Gene Set Enrichment Analysis (GSEA) [32], and KEGG gene sets [33]. Figure 3A retains the archived IPA canonical pathway analysis. Enrichment significance is represented by -log10(*P* value), whereas the activation z-score is an IPA-predicted activity state; these quantities are distinct. Figure 3B-C presents newly recomputed donor-level pre-ranked enrichment using GSEApy v1.3.0 [34] with Hallmark [35] and legacy KEGG definitions from MSigDB v2022.1.Hs. The KEGG file matched the archived gene-set file by SHA-256 checksum.

Ensembl gene IDs were mapped to approved symbols using the HGNC complete set downloaded on 6 October 2026. For repeated symbols, the gene with the highest mean normalized count was retained. Finite Wald statistics for 17,125 unique symbols were ranked, with positive values indicating higher expression in T2D EV-treated recipient cells. Adaptive multilevel sampling used weight 1, sample size 101, eps 1e-10, random seed 20261006, and gene-set sizes 15-500 after intersection with the ranked universe. Fifty Hallmark and 164 KEGG sets were tested. Nominal enrichment *P* values were adjusted separately within each collection using the Benjamini-Hochberg procedure [28]. These tests used a ranked-gene null rather than donor-label permutations.

### Synthetic miRNA mimic transfection and ZEB1 analysis

DU145 cells were transfected using Lipofectamine RNAiMAX (Invitrogen, 13778150). Synthetic mimics were obtained for hsa-miR-192-5p (MC10456), hsa-miR-29b-3p (MC10103), and hsa-miR-106b-3p (MC12321). A working final mimic concentration of 15 nM was reconstructed from the retained laboratory protocol; a negative-control mimic was included. Medium was replaced 24 h after transfection, followed by an additional 48 h of culture before RNA isolation and assessment of *ZEB1* expression by RT-qPCR. *ZEB1* fold-change values were analyzed by one-way ANOVA followed by Tukey’s multiple-comparisons test. Data are presented as mean ± SEM, with n=3 samples per condition.

### Animal handling

Animal experiments were approved by the Boston University Institutional Animal Care and Use Committee. Six-week-old female C57BL/6J mice were obtained from The Jackson Laboratory, acclimated for 1 week, and maintained with ad libitum access to food and water. HFD-fed mice received a diet containing 60% of calories from fat (Research Diets, D12492i); control mice received a matched low-fat diet containing 10% of calories from fat (Research Diets, D12450Ji). Mice remained on these diets for 12 weeks before metabolic assessment. No formal *a priori* power calculation was performed.

### Glucose tolerance testing

Glucose tolerance was assessed after 12 weeks of dietary treatment. Mice were fasted for 6 h, after which baseline tail-vein blood glucose was measured using an AlphaTrak 3 glucometer and compatible test strips. Glucose was administered by oral gavage at 2 g/kg body weight, and blood glucose was measured at 15, 30, 60, and 120 min. Glucose concentrations were plotted over time and area under the curve (AUC) for glucose response was calculated. After assignment of HFD-fed mice to untreated HFD or HFD+metformin groups, glucose tolerance testing was repeated at weeks 4 and 8 of treatment.

### Metformin treatment

After the initial dietary period and glucose-tolerance assessment, HFD-fed mice were assigned to untreated HFD or HFD+metformin groups and remained on HFD. Metformin was supplied in drinking water at 1.8 mg/mL. Glucose was added to improve palatability, with an equivalent amount added to the drinking water of untreated HFD mice. Glucose tolerance was assessed at weeks 4 and 8 of metformin treatment. Plasma from individual mice was subsequently used for EV isolation and RNA experiments.

### Mouse plasma EV-derived RNA isolation and transfection

RNA was isolated from mouse plasma EV-enriched fractions using the miRNeasy Serum/Plasma Kit (Qiagen, cat. no. 217184). Preparations were lysed with QIAzol, followed by chloroform phase separation, ethanol addition, binding to an RNeasy MinElute spin column, washing, and elution in RNase-free water according to the manufacturer’s protocol. DU145 cells were transfected using Lipofectamine RNAiMAX (Invitrogen, cat. no. 13778150). For a mixture prepared for three wells of a 6-well plate, 9 µL of EV-derived RNA was diluted in 450 µL Opti-MEM, and 27 µL RNAiMAX was separately diluted in 450 µL Opti-MEM. Equal volumes (450 µL each) of the diluted RNA and RNAiMAX solutions were combined and incubated for 5 min, and 250 µL of the resulting transfection mixture was added to each well. Medium was replaced after 24 h, followed by an additional 48 h of culture before functional assessment. The negative transfection control was mirVana™ miRNA Mimic Negative Control (Invitrogen/Thermo Fisher Scientific, cat. no. 4464058). EV-derived RNA derived from individual mice was maintained separately throughout the experiment and was not pooled.

### Transwell migration and Matrigel invasion assays

After transfection with plasma EV-derived RNA, DU145 cells were maintained for 72 h, with medium replacement after the first 24 h, as described above. Cells were then serum-starved in serum-free DMEM for 3 h, trypsinized, and seeded into the upper compartments of 24-well Transwell inserts with 8-µm pores. Migration was assessed using uncoated inserts, whereas invasion was assessed using Corning BioCoat Matrigel Invasion Chambers with 8.0-µm PET membranes (Corning, cat. no. 354480), with 75,000 cells seeded per invasion or migration insert. Matrigel chambers were rehydrated according to the manufacturer’s instructions before use. DMEM containing 10% FBS was placed in the lower chamber as a chemoattractant. Assays were incubated for 24 h at 37 °C in 5% CO□.

After incubation, cells remaining on the upper membrane surface were removed with a cotton swab. Cells on the lower surface were fixed in absolute methanol for 5 min at −20 °C, stained with 1% crystal violet in 2% ethanol for 10 min, washed with DPBS, and air-dried. Membranes were imaged using an EVOS XL Core digital inverted microscope. Migration and invasion were quantified in Fiji/ImageJ as the percentage of analyzed membrane area occupied by crystal-violet-positive staining [36]. Local background was corrected, and staining was quantified within the interior membrane region after exclusion of edge and insert-rim artifacts.

Three independent HFD mice and three independent HFD+metformin mice were analyzed; technical measurements were not counted as independent biological replicates. EV-derived RNA from each mouse was maintained separately, and three technical insert/image-set measurements per mouse were averaged to generate one mouse-level value. Treatment comparisons therefore used the mouse as the biological experimental unit (n=3 mice per group).

### Statistical analysis

Nominal *P* values from the 84-miRNA plasma EV screening panel were additionally adjusted using the Benjamini-Hochberg procedure [28] across all assayed miRNAs. RNA-seq differential expression was evaluated using adjusted *P* values as described above. For Figure 3B-C, nominal enrichment *P* values were adjusted separately within the Hallmark and KEGG collections using the Benjamini-Hochberg procedure. ZEB1 fold-change values after miRNA mimic transfection were analyzed using one-way ANOVA followed by Tukey’s multiple-comparisons test. For the migration experiment, the individual mouse was the biological experimental unit (n=3 mice per group). Image measurements were averaged within each mouse, and the resulting mouse-level values were compared using a two-sided Welch t-test. For the invasion assay, three technical area-coverage measurements from each mouse were averaged with equal weight, and the resulting values from three independent mice per treatment were compared using an unpaired, two-sided Welch t-test. The migration and invasion cohorts were analyzed separately. Scramble controls were displayed descriptively and were not included in the mouse-level treatment comparison. Statistical significance was defined as *P*<0.05 unless otherwise specified.

## Results

### Plasma EV miRNA profiles show differences between T2D and non-diabetic donors

To determine whether the miRNA composition of plasma EV-enriched fractions differed according to metabolic status, we profiled 84 cancer-associated miRNAs in EV-derived RNA from six independent T2D donors and six independent ND donors. Several miRNAs showed nominal differences in abundance between groups; however, none remained statistically significant after Benjamini-Hochberg correction across the 84-miRNA panel. Accordingly, the miRNA profiling experiment was considered exploratory (Fig. 1A-B). miR-192-5p, miR-106b-5p, and miR-29b-3p each showed greater than two-fold elevation in the T2D group together with nominal *P*<0.05 and were carried forward as candidate signals rather than as an FDR-validated T2D EV-miRNA signature.

DU145 cells were transfected with synthetic mimics of hsa-miR-192-5p, hsa-miR-29b-3p, and hsa-miR-106b-3p. The predefined one-way ANOVA/Tukey analysis identified significant increases in ZEB1 expression relative to the negative-control mimic for miR-106b-3p and miR-192-5p (Tukey-adjusted *P*=0.0022 and *P*=0.00015, respectively), whereas miR-29b-3p did not significantly alter *ZEB1* expression (Tukey-adjusted *P*=0.34; Fig. 1C).

### Transcriptional responses of DU145 cells to plasma EVs from T2D and non-diabetic donors

To examine the global transcriptional response of prostate cancer cells to circulating EVs, DU145 cells were treated with intact plasma EV-enriched fractions from six independent T2D donors and two independent ND donors, followed by RNA sequencing after three days of treatment. For selected donors, multiple recipient-cell treatment wells were sequenced; these represent within-donor recipient-cell replicates rather than additional independent human donors.

In the PCA visualization, untreated control and ND EV-treated recipient-cell samples occupied similar regions of PCA space, whereas T2D EV-treated samples showed greater dispersion (Fig. 2A). These patterns describe variation across donor exposures, with multiple recipient-cell treatment wells analyzed for each donor; the contribution of individual medications was not assessed. In the differential-expression analysis used to generate the T2D versus ND volcano plot, 2,984 genes met adjusted *P*<0.05 and |log2FC|>1 (Fig. 2B). The Figure 2C heatmap used the subset of genes meeting these criteria in both the T2D versus ND and T2D versus untreated control comparisons.

### Exposure to T2D plasma EV-enriched fractions is associated with cytoskeletal and extracellular-matrix pathway enrichment

Pathway analyses identified differences in gene-expression patterns related to extracellular matrix organization, cytoskeletal regulation, mitosis and RHO-GTPase signaling following exposure to T2D plasma EV-enriched fractions (Fig. 3A). These findings highlighted processes involved in cell structure, interactions with the surrounding matrix and cell division.

To examine these patterns at the donor level, we compared recipient-cell transcriptional responses to EV-enriched fractions from six T2D and two ND donors. Among the displayed Hallmark gene sets, mitotic spindle was enriched toward T2D, whereas MYC targets V1 were enriched toward ND (Fig. 3B). Within the KEGG collection, ECM–receptor interaction was enriched toward T2D, while cardiac muscle contraction and ribosome were enriched toward ND (Fig. 3C). All five displayed gene sets had BH-adjusted enrichment P values below 0.05 within their respective tested collections. Together, these analyses identify distinct transcriptional responses to T2D and ND plasma EV-enriched fractions, with enrichment of mitotic spindle and ECM–receptor interaction genes toward T2D.

### Systemic metformin treatment is associated with reduced pro-migratory activity of plasma EV-derived RNA

After 12 weeks of HFD feeding, glucose tolerance was assessed before assignment to HFD or HFD+metformin treatment groups. Glucose tolerance was subsequently assessed at weeks 4 and 8 of metformin treatment. Glucose-response profiles were similar between the HFD and HFD+metformin groups at week 4, whereas by week 8 the HFD+metformin group showed a lower glucose response consistent with improved glucose tolerance (Fig. 4A).

**Figure 4.**
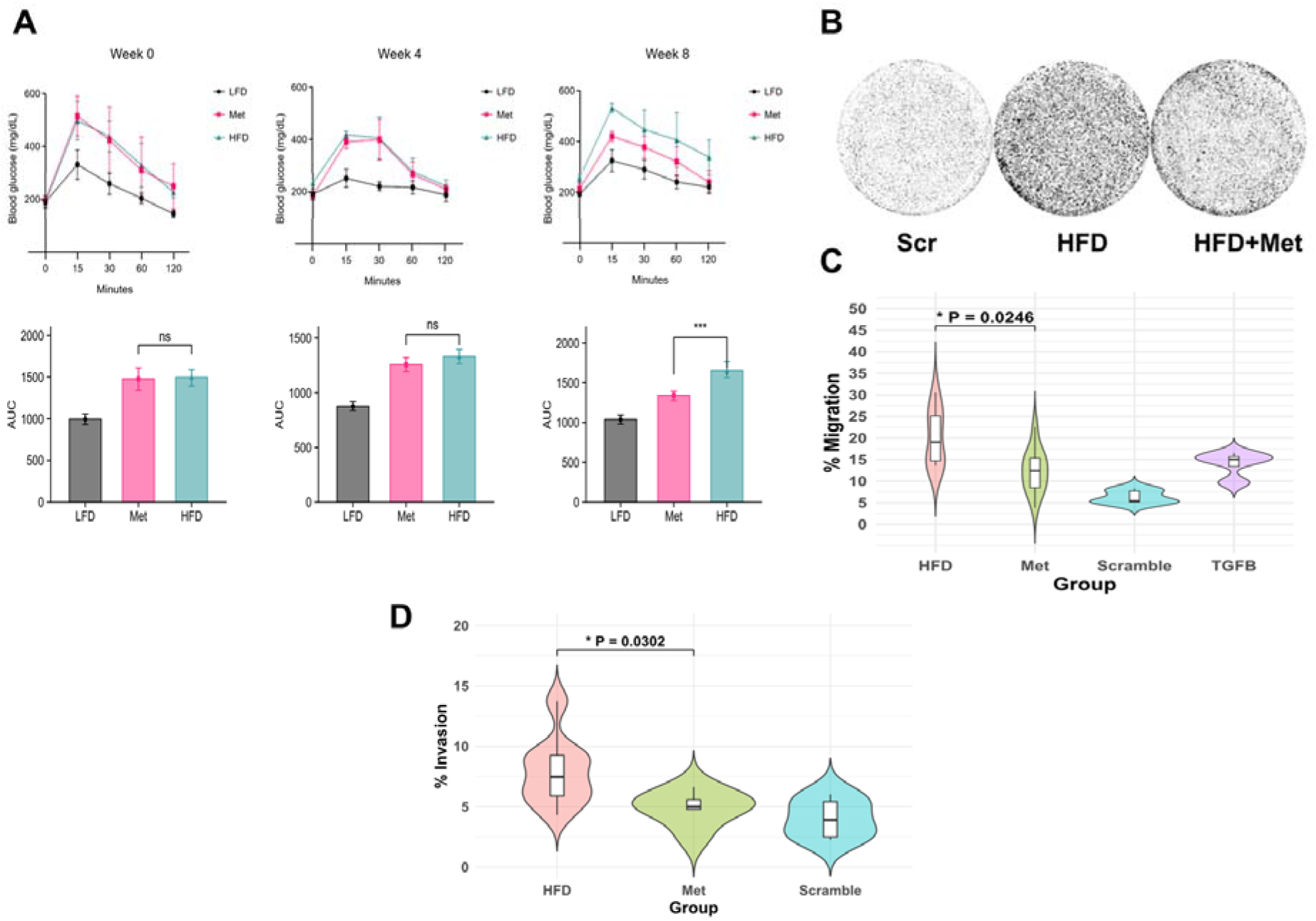
Systemic metformin treatment and the functional activity of plasma EV-derived RNA in DU145 cells. A. Glucose tolerance testing and corresponding AUC measurements during the HFD/metformin treatment study. HFD+metformin mice remained on HFD during treatment. B. Representative images from transwell migration assays after transfection of DU145 cells with scramble (Scr) control or plasma EV-derived RNA isolated from individual HFD or HFD+metformin mice. C. Violin distributions and internal boxplots of well/image-level migration measurements. For HFD and Met (HFD+metformin; n=3 independent mice/group). The bracket reports the mouse-level two-sided Welch t-test, \**P*=0.0246. Scramble and TGFB controls are shown descriptively. D. Violin distributions and internal boxplots of well/image-level invasion measurements. For HFD, Met (HFD+metformin; n=3 independent mice/group) and Scramble control. Three technical measurements were averaged within each mouse (n=3 mice/treatment) besides scrambled control. The bracket reports the unpaired, two-sided mouse-level Welch t-test, \**P*=0.0302.

Plasma EV-derived RNA was isolated separately from individual HFD and HFD+metformin mice and transfected into DU145 cells. DU145 cells receiving EV-derived RNA from HFD+metformin mice showed lower migration than cells receiving EV-derived RNA from untreated HFD mice (mean membrane coverage, 12.23% versus 20.25%; n=3 mice per group; two-sided Welch t-test, *P*=0.0246; Fig. 4B-C).

### Matrigel invasion measurements after treatment with mouse plasma EV-derived RNA

In addition to migration, we assessed invasive behavior. DU145 cells were transfected with plasma EV-derived RNA from an independent cohort of HFD or HFD+metformin mice and evaluated in a Matrigel invasion assay. Figure 4D shows nine technical insert/image-set measurements per group. Three measurements were averaged within each mouse before comparing three mice per treatment. Mean stained membrane coverage, averaged for each mouse, was 7.89 ± 1.20% in the HFD group and 4.89 ± 0.98% in the HFD+metformin group (mean ± SD). Coverage was therefore 3.00 percentage points lower in the HFD+metformin group (95% confidence interval, 0.48–5.52; unpaired, two-sided Welch’s t-test, *P* = 0.0302). Mean coverage in the scramble-control group was 4.04%.

## Discussion

Type 2 diabetes (T2D) has a complex relationship with prostate cancer. Reported inverse associations with incidence vary by population and disease stage: a recent meta-analysis found lower early-stage incidence but no clear association for all stages combined or advanced disease [7], whereas other meta-analytic and cohort studies have reported inverse associations in some populations [8,9]. After diagnosis, some studies have associated diabetes with higher mortality [10] or worse treatment-related outcomes [11–13]. Higher preoperative HbA1c has been associated with subsequent metastases and castration-resistant progression after radical prostatectomy [14], and diabetic-range fasting glucose after diagnosis with higher prostate cancer-specific mortality [15]. Other cohorts report different patterns, including lower prostate cancer-specific mortality at higher HbA1c levels in veterans with T2D [16] and treatment-specific survival differences by BMI and glycemic status [17]. Evidence linking cardiometabolic disease to the risk of developing aggressive prostate cancer [18] should be distinguished from these post-diagnosis findings. Associations with biochemical recurrence remain less consistent [19]. These observations emphasize the importance of separating prostate cancer incidence from post-diagnosis disease behavior. The present study addresses the latter question by examining whether T2D-associated systemic metabolic state is accompanied by altered circulating EV signaling to well established cellular models of human prostate cancer.

Profiling of 84 cancer-associated miRNAs in plasma EV-enriched fractions identified several candidates that differed between T2D and ND donors. MiR-192-5p, miR-106b-5p, and miR-29b-3p each showed greater than two-fold elevation in the T2D group together with nominal screening P<0.05. Among these candidates, miR-192-5p is of particular interest because circulating miR-192 has been linked previously with diabetes, obesity-associated inflammation, and metabolic dysfunction [37–39], and prior work from our group identified increased miR-192-5p in plasma EV preparations from insulin-resistant mice with diet-induced obesity [25]. These observations provide biological context for miR-192-5p but do not establish a T2D-specific plasma EV miRNA signature in the present human cohort.

To examine whether selected candidate miRNAs promote cancer-relevant transcription in recipient cells, DU145 cells were transfected with synthetic mimics of miR-192-5p, miR-29b-3p, and miR-106b-3p. MiR-192-5p and miR-106b-3p increased *ZEB1* expression relative to the negative-control mimic, whereas miR-29b-3p did not significantly increase *ZEB1* expression. *ZEB1* is an established regulator of EMT in prostate cancer [40], and clinical tissue studies have associated EMT-marker expression, including *ZEB1*, with more advanced prostate cancer states [41]. Experimental *ZEB1* knockdown has also been shown to reduce prostate cancer stem-cell-like properties, further supporting its relevance to aggressive cellular phenotypes [42]. Nevertheless, increased *ZEB1* expression in DU145 cells alone does not establish a complete EMT program, increased invasion, metastasis, or clinical progression. The absence of a significant *ZEB1* response to miR-29b-3p should likewise not be interpreted as evidence that miR-29b is biologically inactive; prior prostate cancer work has implicated miR-29b in regulation of invasive behavior [43]. An additional limitation is that the plasma EV profiling panel measured miR-106b-5p, whereas the synthetic mimic experiment used miR-106b-3p. The latter therefore represents an independent functional observation rather than direct validation of the screening signal. Quantitative studies have reported low average miRNA copy numbers per vesicle and inefficient functional delivery in some model systems [44,45]. These findings reinforce the need to distinguish synthetic mimic responses from physiological EV-mediated RNA transfer.

The synthetic miRNA experiments complement the EV studies by demonstrating that selected RNA species can directly modify a cancer-relevant transcriptional program in recipient cells. These experiments are necessarily reductionist and therefore provide mechanistic information distinct from that obtained with intact EV preparations, which deliver a complex combination of RNAs, proteins, lipids, and other cargo. NTA provided donor-matched measurements of particle size and concentration for the human plasma EV-enriched preparations. We therefore use the operational term EV-enriched fractions, while future studies incorporating orthogonal EV-characterization approaches and direct cargo profiling can build on these measurements to define particle identity in greater detail and identify the molecular components responsible for the observed biological effects [20,55].

To assess the broader effects of intact circulating EV preparations, DU145 cells were treated with plasma EVs purified from T2D or ND donors and subjected to RNA sequencing. Exposure to T2D-derived preparations was associated with broad transcriptional differences relative to ND-derived preparations, together with substantial heterogeneity among T2D EV-treated recipient cells. The heterogeneity observed among T2D donor-derived EV treatments may reflect multiple donor-level factors, including metabolic status, disease duration, medication exposure, and other clinical characteristics. Although medication history originally suggested a possible relationship between metformin exposure and the transcriptional patterns observed in the human RNA-sequencing experiment, the cohort was not designed or powered to isolate medication effects, and no formal analysis demonstrated a metformin-specific association. The human data should therefore be considered hypothesis-generating with respect to medication exposure.

The pathway analyses revealed distinct transcriptional responses to plasma EVs from T2D and ND donors. In the donor-level analysis, genes involved in mitotic spindle organization and extracellular matrix–receptor interactions were preferentially enriched in DU145 cells treated with T2D-derived EVs. In contrast, MYC target genes, ribosomal pathways, and cardiac muscle contraction gene sets showed enrichment toward the ND-treated group (Fig. 3B-C). The complementary IPA analysis also identified RHO-GTPase and cytoskeletal signaling among the prominent pathways associated with T2D EV exposure (Fig. 3A). The prominence of RHO-family pathways is particularly notable, because these small GTPases are central regulators of actin remodeling, cell polarity, adhesion, and motility, suggesting a potential mechanistic connection between the transcriptional changes observed here and the migratory phenotypes associated with metabolically altered EV signaling. Previously, our group reported a closely related pattern in an independent breast cancer model, in which plasma EVs from HFD-fed, insulin-resistant mice increased migration and metastatic behavior of E0771 triple-negative breast cancer cells and were associated with enrichment of RHO-GTPase signaling, particularly Rac1 [25]. Thus, the recurrence of RHO-GTPase-associated signaling across metabolically altered EV exposures in both prostate and breast cancer models suggests that cytoskeletal reprogramming may represent a conserved downstream response to systemic metabolic dysfunction. Consistent with this interpretation, prior work from our group showed that purification of EV RNA by acidic phenol extraction and isopropanol precipitation enhanced the transcriptional effects observed in recipient tumor cells [26], supporting EV-derived RNA as an important bioactive component of this signaling axis. Together, these findings connect metabolic dysfunction, EV-derived RNA signaling, RHO-family GTPase pathways, and cytoskeletal remodeling across independent experimental systems and provide a mechanistic framework for investigating the altered migratory and invasive behavior observed in the present study.

To examine metformin under controlled conditions, we used HFD-fed mice treated with or without systemic metformin and assessed the effects of RNA isolated from their plasma EV-enriched fractions on DU145 cell behavior. EV-derived RNA from HFD+metformin mice induced less DU145 migration than RNA from untreated HFD mice. Importantly, this finding was complemented by an independent Matrigel invasion experiment, which also showed lower mean stained membrane coverage in the HFD+metformin group (n=3 mice/group; Welch P=0.0302). Although each experiment included three mice per treatment group, the concordant reduction in migration and invasion across independent cohorts strengthens the biological significance of these observations. Together, these findings support a metformin-associated difference in the functional activity of circulating EV-derived RNA and raise the possibility that systemic metformin treatment may attenuate aggressive behavior of prostate cancer cells through an additional, indirect signaling mechanism.

The functional differences observed with EV-derived RNA from metformin-treated mice are consistent with a broader literature showing that systemic metformin treatment can reprogram prostate cancer biology through several potential routes. Experimental studies have reported direct effects of metformin on prostate cancer-cell migration and signaling [46,47], and broader reviews have summarized potential antitumor mechanisms and clinical applications of metformin in prostate cancer [48]. Outside prostate cancer, metformin has also been shown to alter the extracellular secretory environment; for example, AMPKα1-dependent suppression of extracellular Hsp90α secretion reduced tumor-cell migration, invasion, and metastasis in experimental models [49]. Together, these findings provide mechanistic precedent for the ability of metformin to modify extracellular signaling and attenuate tumor-cell aggressive behavior. In the context of the present study, this raises the intriguing possibility that changes in circulating EV-derived signals represent an additional route through which systemic metformin treatment may reduce prostate cancer-cell migration and invasion.

At the clinical level, a growing body of observational and real-world evidence has examined metformin exposure in relation to prostate cancer incidence, recurrence, and mortality, suggesting that its effects may depend on disease stage, treatment context, and the metabolic characteristics of the patient population [50,51]. Randomized trials have likewise highlighted this complexity. In TAXOMET, addition of metformin to docetaxel did not improve prostate-specific antigen response, progression-free survival, or overall survival in metastatic castration-resistant prostate cancer [52], while the phase III MAST trial did not demonstrate reduced progression among men with low-risk prostate cancer managed with active surveillance [53]. More recently, STAMPEDE found that adding metformin to standard-of-care therapy for non-diabetic men with metastatic hormone-sensitive prostate cancer did not significantly improve overall survival in the overall study population but importantly reduced several adverse metabolic consequences of androgen-deprivation therapy [54]. Together, these findings suggest that the clinical effects of metformin in prostate cancer are context dependent, emphasizing the importance of understanding its biological actions beyond direct effects on tumor cells. The present study contributes to this emerging perspective by identifying changes in the functional activity of circulating EV-derived RNA as a potential additional consequence of systemic metformin treatment, providing a mechanistic direction for future investigation.

Building on these findings, further investigation of circulating EV signaling may reveal important molecular mechanisms linking systemic metabolic dysfunction with prostate cancer-cell behavior. In the human cohort, validation of the candidate miRNAs identified in the plasma EV screen in larger, independently characterized populations will help establish their biological relevance. Furthermore, the inter-individual variability observed in the RNA-sequencing experiments highlights the importance of examining how donor-specific metabolic characteristics and medication exposure influence EV-mediated communication with tumor cells. In the mouse model, an important next step will be to identify the specific RNA species responsible for the observed reductions in migration and invasion. Comparative molecular profiling of EV-derived RNA from HFD and HFD+metformin mice, followed by functional validation of candidate miRNAs and other RNA species, may clarify how systemic metformin treatment modifies the biological activity of circulating EV-enriched fractions. Additional studies examining EV cargo composition, abundance, release, clearance, and tissue origin will further advance our understanding of these processes. Integrating these approaches with detailed clinical annotation and metabolic phenotyping may ultimately help distinguish shared metabolic signatures from donor-specific responses and identify molecular pathways through which systemic metabolic interventions reprogram tumor-cell behavior.

Taken together, our findings support a model in which systemic metabolic dysfunction creates changes in circulating EV signaling that drive adverse prostate cancer-cell responses. Human plasma EV-enriched fractions from donors with T2D exhibited differences in candidate miRNA abundance and induced distinct transcriptional responses in DU145 cells, highlighting the potential of circulating EV-derived signals to reflect systemic metabolic alterations. Importantly, experiments using RNA isolated from plasma EV-enriched fractions of metformin-treated HFD mice demonstrated reduced DU145 migration and invasion in two independent mouse cohorts, providing complementary functional evidence that systemic metformin treatment reduces the tumor progression-associated biological activity of RNA contained in circulating EVs. Although the specific RNA mediators and underlying molecular mechanisms remain to be identified, these findings establish a foundation for investigating circulating EV-derived RNA as a link between systemic metabolic state and prostate cancer-cell behavior. More broadly, this work highlights extracellular vesicle communication as a promising avenue for understanding how metabolic dysfunction and metabolic interventions may promote cancer progression.

## Data availability

Raw and processed RNA-sequencing data have been deposited in the NCBI Gene Expression Omnibus (GEO) under accession GSE349995 (https://www.ncbi.nlm.nih.gov/geo/query/acc.cgi?acc=GSE349995). The dataset is currently held private pending publication. Analysis files and other data supporting the findings of this study are available from the corresponding author upon reasonable request.

## Author contributions

Conception and design: M.S., G.V.D. Methodology: M.S., P.L., Y.Q., C.S.E., A.C., S.M., G.V.D. Acquisition of data: M.S., P.L., H.K., Y.Q., C.S.E. Clinical coordination: H.K., S.A., D.S. Analysis and interpretation of data: M.S., A.C. Writing, editing, and revision of manuscript: M.S., P.L., G.V.D. Study supervision and funding: G.V.D.

## Acknowledgements

We thank Dr. Carla Mazzeo for expert opinion and consulting about extracellular vesicle analysis and quality control; she was co-author of MISEV2023, the international consensus guidelines for EV research, and here helped us use analytical rigor and pre-analytical variable control required to generate reproducible, clinically translatable exosome biomarker data. We also thank Dr. Yuriy Alekseyev of the Microarray and Sequencing Resource at BUMC for technical assistance. This work was supported by the Cancer Moonshot and Cancer Systems Biology Consortium of the National Cancer Institute (G.V.D.: U01CA182898, U01CA243004, and R01CA222170), as well as the Shipley Prostate Cancer Research Center and the Chung-Ho Chen Cancer Research Fund at the Chobanian & Avedisian School of Medicine (GVD).

