## Supplementary figures and images for "Metabolic dysfunction and metformin modify plasma extracellular vesicle signaling to prostate cancer cells"

### Supplementary Fig S1

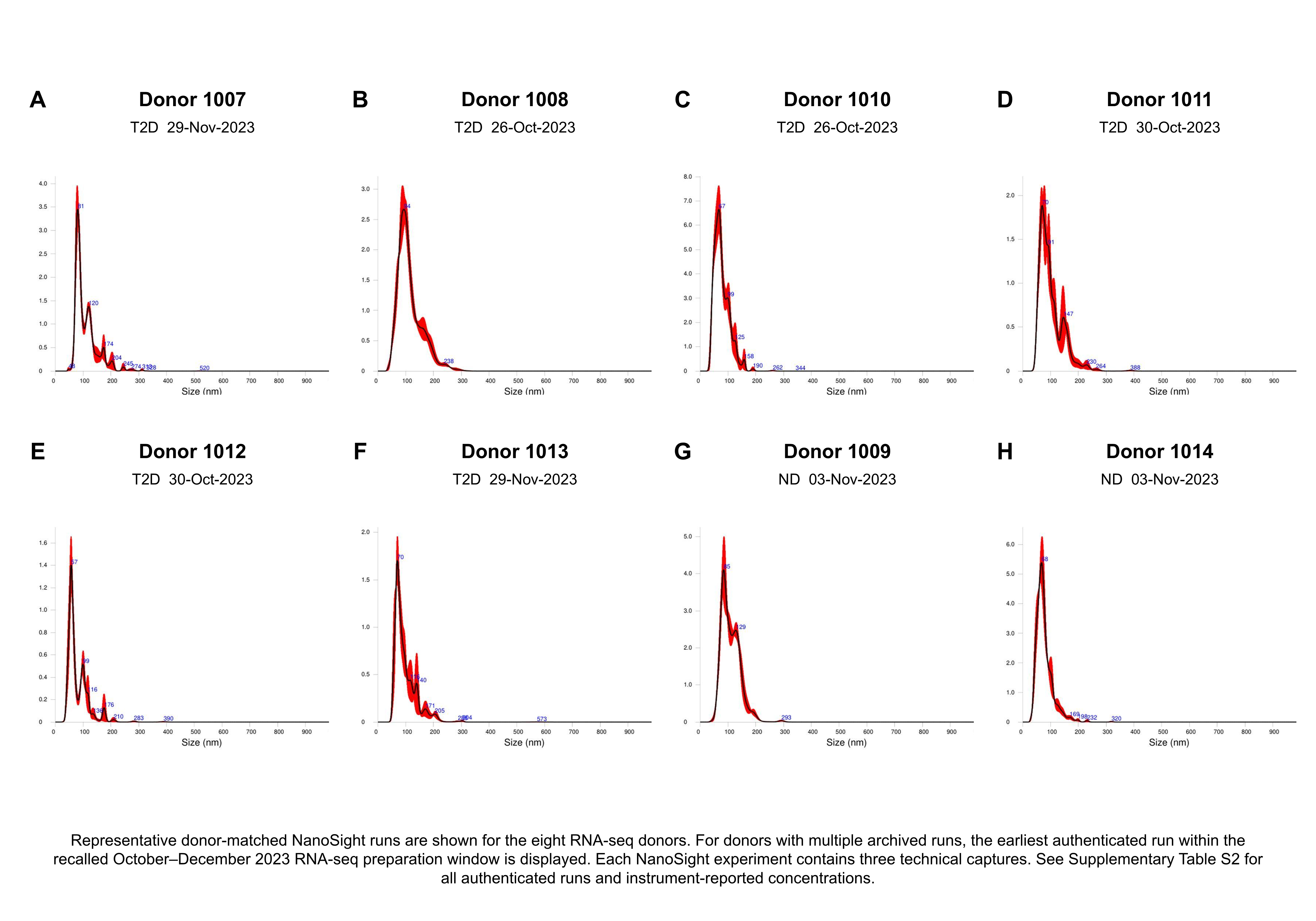

### Supplementary Table S1

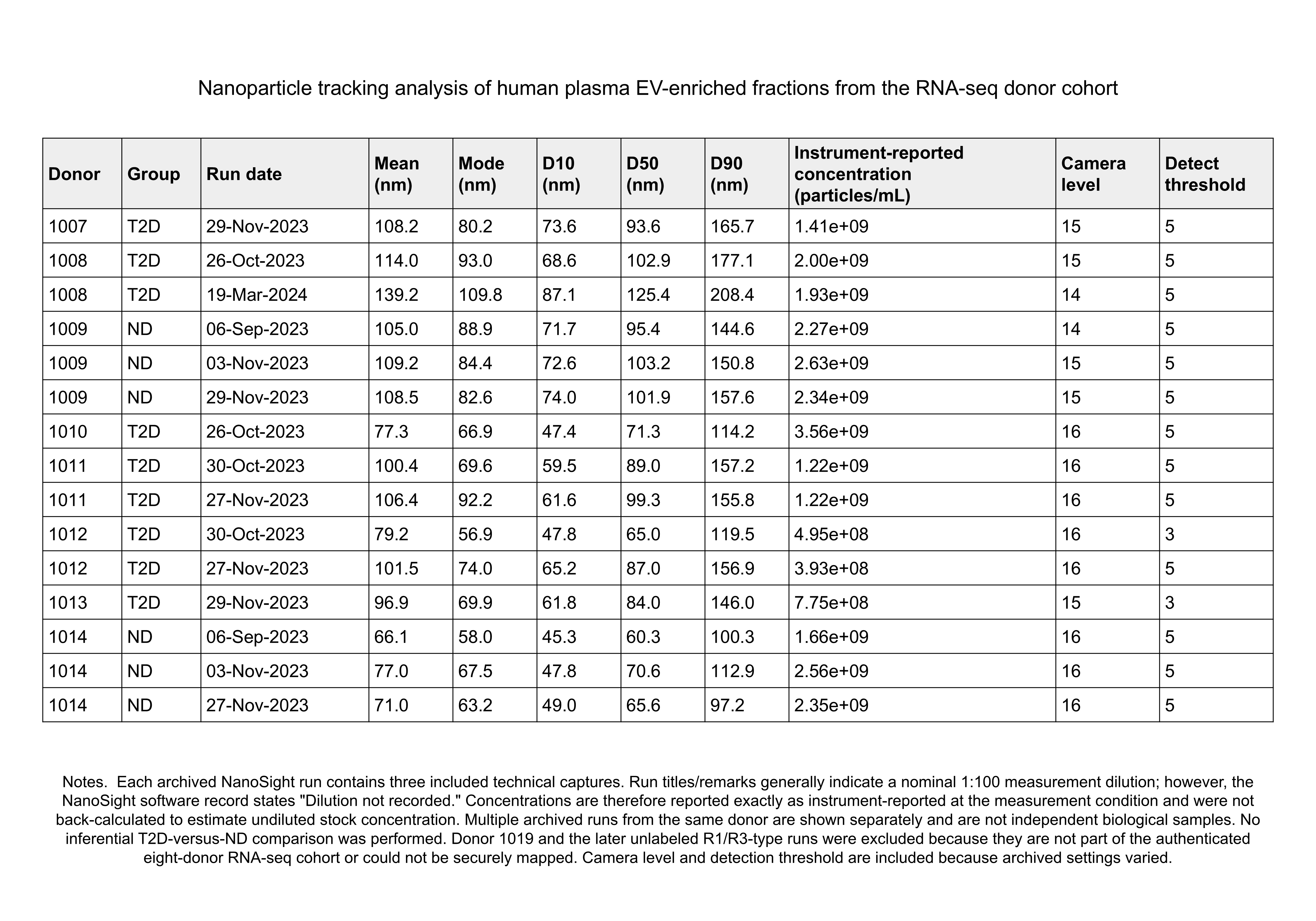

### Table 1

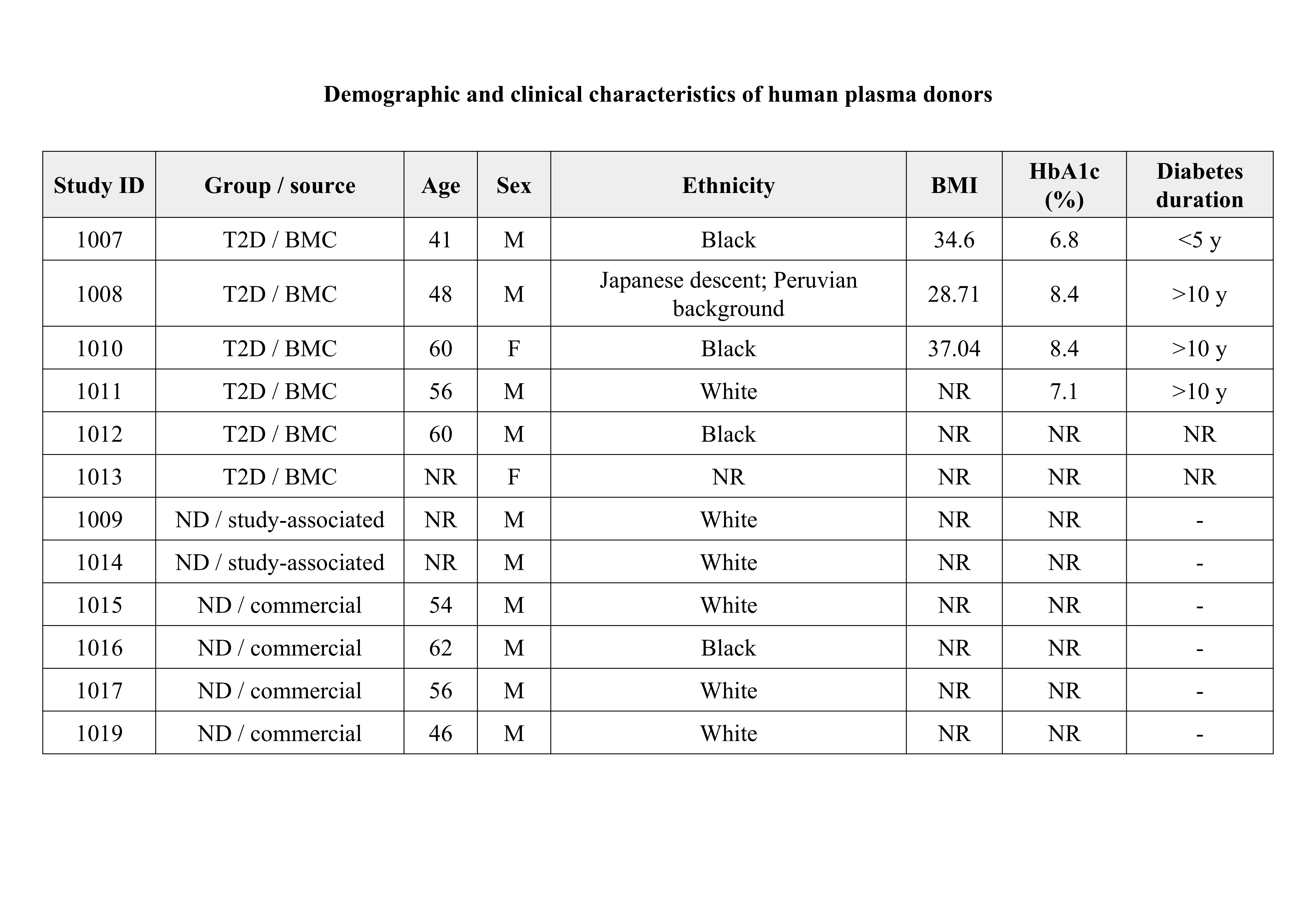

### Table 2

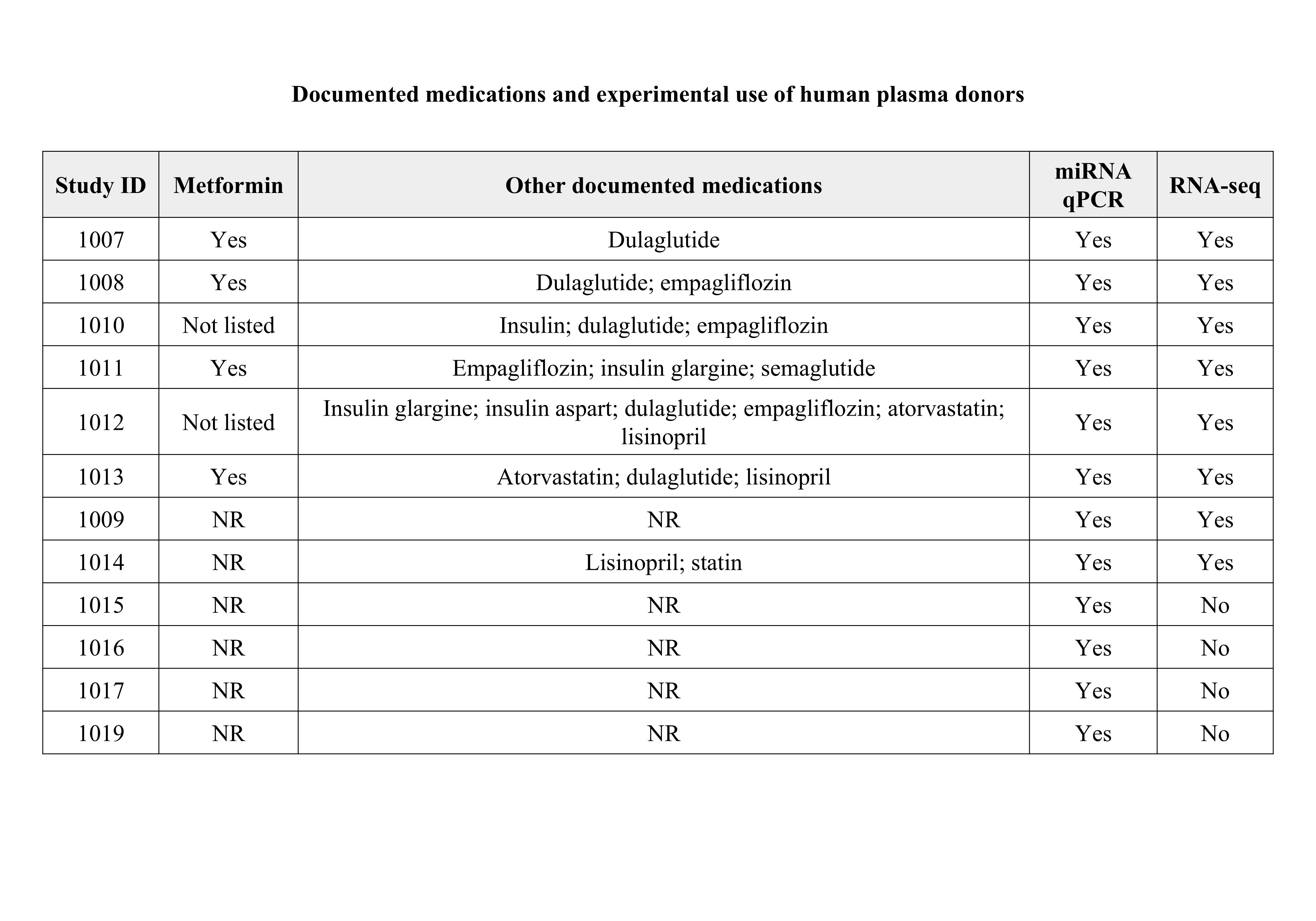
