## Supplementary Figure legends for "Metabolic dysfunction and metformin modify plasma extracellular vesicle signaling to prostate cancer cells"

**Supplementary Figure S1. Nanoparticle tracking analysis of human plasma EV-enriched fractions.** Representative NanoSight NS300 particle-size distributions are shown for the eight human donors included in the RNA-sequencing cohort: six donors with type 2 diabetes (T2D) and two non-diabetic (ND) donors. Panels A–F show T2D donors 1007, 1008, 1010, 1011, 1012, and 1013, respectively; panels G–H show ND donors 1009 and 1014. Donor identifiers and measurement dates are shown above each profile. Each NanoSight run includes three technical captures. Particle-size measurements, instrument-reported concentrations, and acquisition settings for all included runs are provided in Supplementary Table S1. EV, extracellular vesicle.

**Supplementary Table S1. Particle-size measurements and instrument-reported concentrations of human plasma EV-enriched fractions.** Fifteen archived NanoSight NS300 runs are listed for the eight RNA-sequencing donors shown in Supplementary Figure S1. Columns report donor identifier, group, run date, mean and modal particle diameter, D10, D50, D90, instrument-reported concentration, camera level, and detection threshold. Diameters are expressed in nanometers; D10, D50, and D90 denote the 10th, 50th, and 90th percentiles of the particle-size distribution, respectively. Concentrations are expressed as particles/mL. Each run includes three technical captures. Camera level and detection threshold are included because acquisition settings varied between runs. T2D, type 2 diabetes; ND, non-diabetic; EV, extracellular vesicle.
